# Rapid species discovery and identification with real-time barcoding facilitated by ONTbarcoder 2.0 and Oxford Nanopore R10.4

**DOI:** 10.1101/2023.06.26.546538

**Authors:** Amrita Srivathsan, Vivian Feng, Daniel Suárez, Brent Emerson, Rudolf Meier

**Affiliations:** Center for Integrative Biodiversity Discovery, Leibniz Institute for Evolution and Biodiversity Science, Museum für Naturkunde, Invalidenstraße 43, 10115 Berlin, Germany; Island Ecology and Evolution Research Group, Institute of Natural Products and Agrobiology (IPNA-CSIC), C/Astrofísico Francisco Sánchez 3, La Laguna, Tenerife, Canary Islands 38206, Spain; School of Doctoral and Postgraduate Studies, University of La Laguna, 38200 La Laguna, Tenerife, Canary Islands 38200, Spain; Department of Biological Sciences, National University of Singapore, 14 Science Drive 4, Singapore, 117543, Singapore

## Abstract

Most arthropod species are undescribed and hidden in biodiversity samples that are difficult to sort to species using morphological characters. Sorting specimens to putative species with DNA barcodes is an attractive alternative, but needs cost-effective techniques that are suitable for use in many laboratories around the world. Barcoding with Oxford Nanopore’s portable and inexpensive MinION sequencers could be a good presorting approach because it reduces the space and capital cost of a fully functional barcoding laboratory. Similarly important would be user-friendly and reliable software for analysis of the ONT data. It is here provided in the form of an updated ONTbarcoder that is available for all commonly used operating systems and includes a GUI. The new version of ONTbarcoder has three key improvements related to the higher read quality obtained with ONT’s latest basecallers and chemistry (R10.4 flow cells, Q20+ kits). Firstly, ONTbarcoder can now deliver real-time barcoding to complement ONT’s real-time sequencing. This means that the first barcodes are obtained within minutes of starting a sequencing run and a significant proportion of barcodes are available before sequencing completes. This real-time feature of ONTbarcoder can be configured to handle both Mk1B and Mk1C sequencers. The input is a demultiplexing sheet and sequencing data (raw or basecalled). Secondly, the improved read quality of ONT’s latest flow cells (R10.4) allows for the use of primers with shorter indices than those previously needed (9 bp vs. 12-13 bp). This decreases primer cost and improves PCR success rates. Thirdly, we demonstrate that the availability of R10.4 chemistry in the low-cost Flongle flow cell is an attractive option for users who require only 200-250 barcodes at a time.

## Introduction

The world faces significant challenges as we head into a biodiversity crisis affecting many ecosystems. At the heart of this crisis is the loss of species, habitats, and ecological interactions that have evolved over millions of years (Biesmeijer *et al*., 2006; Outhwaite *et al*., 2022). Monitoring biodiversity loss will be important but very difficult given that over 80% of the animal species are unknown, undescribed and unidentifiable (Oliver *et al*., 2015; Schmeller *et al*., 2020; World Economic Forum, 2020; Srivathsan *et al*., 2023). This problem is particularly acute for arthropods, which account for the vast majority of animal species and specimens. In fact, the biomass of all arthropods combined surpasses that of all terrestrial vertebrates in the wild (Bar-On *et al*., 2018). Arthropods are essential and provide many important ecosystem services including pollination, pest control, nutrient cycling or as food resources (Belovsky and Slade, 2000; Bianchi *et al*., 2006; Gallai *et al*., 2009; Tallamy and Shriver, 2021; Fürst *et al*., 2023). Yet, despite their importance, most species are unknown which creates a fundamental impediment to understanding of biodiversity, whether it be in the context of quantitative biodiversity assessment, understanding of the tree of life, or establishing the role of arthropods in delivering ecosystem functions. The problems with understanding arthropod diversity is in part due to typically high abundance obtained from bulk sampling techniques such as Malaise trapping, which makes it expensive or even impossible to sort the thousands of specimens from bulk samples to species.

Thus, new approaches are needed that allow for the discovery of and taxonomic research on unknown species hidden in bulk samples, together with already describes species. Here, DNA barcodes are a useful tool for identifying and sorting specimens to approximate species level (Wang *et al*., 2018) but until recently, high cost and time-consuming processes for obtaining barcodes have limited its usefulness (Hebert *et al*., 2003; Meier *et al*., 2016; Srivathsan *et al*., 2021).

Fortunately, the advent of simplified methods for obtaining and sequencing amplicons with new sequencing technologies, such as the MinION sequencer from Oxford Nanopore Technologies (ONT), are making DNA barcoding more accessible and cost-effective (Srivathsan *et al*., 2018; Pomerantz *et al*., 2022). These recent developments enable large-scale species discovery and specimen identification at a fraction of the cost of previously available methods. Large-scale barcoding or megabarcoding not only expedites the specimen-sorting step of species discovery, but also improves the data obtained with metabarcoding by providing the barcode reference databases that are very useful for interpreting metabarcoding data (Chua *et al*., 2023). It is therefore important to have cost-effective barcoding protocols that are sufficiently simple to be implemented by stakeholders anywhere within a range of scientific, financial, and skill backgrounds (Srivathsan *et al*., 2021).

We recently proposed such a workflow that covered the three main steps of DNA barcoding (Srivathsan *et al*., 2021). (1) obtaining COI amplicons for a large number of specimens, (2) sequencing a pool of COI amplicons, and (3) generating barcodes from the MinION sequencers. The workflow reduced the cost of a fully functional DNA barcoding laboratory by more than 90% to approximately USD 10,000. Specimen barcodes can be obtained using this workflow at the cost of <0.10 USD per barcode (Srivathsan *et al*., 2021) which is much lower than those obtained using Sanger sequencing or via sequencing facilities (Canadian Center for DNA barcoding: 3-5 CAD, https://ccdb.ca/pricing/). We furthermore introduced a barcoding software (“ONTbarcoder”) that is user-friendly and compatible with all popular operating systems (iOS, Windows, Linux). We here provide an update to the software that responds to improved chemistry and basecalling models released by Oxford Nanopore Technologies. Higher read quality derived from these improvements means that (1) real time sequencing can now be matched by real-time barcoding, (2) the length of primer indices can be reduced, (3) many error correction options in ONTbarcoder are now rarely needed, and (4) all these benefits are now also available for the Flongle, the cheapest flow cell from ONT.

The first novel feature of the latest ONTbarcoder is real-time barcode calling to match Oxford Nanopore’s real-time sequencing. ONT devices are unique in that they produce full-length reads within minutes of starting a sequencing run. The problem with matching real-time sequencing with real-time barcode calling has been the high error rates of ONT reads. All earlier attempts of implementing real-time barcoding yielded unsatisfying results due to low demultiplexing and high sequence error rates (Srivathsan *et al*., 2018, 2019). All analysis was thus relegated to after all reads had been generated. The second novel feature of the barcoding pipeline proposed here is the use of shorter primer tags (“indices”). Previously, the tags needed to be 12-13 base pairs (bp) long (Srivathsan *et al*., 2018, 2019). This increased primer cost and interfered with the success rate of PCRs (Srivathsan *et al*., 2019; Bohmann *et al*., 2022), because a significant proportion of the primer (i.e., the tag) did not match the template. We here show that the tag length can now be reduced to 9 bp. Higher read quality also underpins the third improvement, i.e., all three levels of the iterative pipeline in ONTbarcoder (“Consensus by length”, “Consensus by similarity”, “Consensus by barcode comparison”) are rarely needed. This leads to an improvement in speed as well as reduction in number of ambiguities in resulting barcodes. Lastly, all the benefits of the new chemistry are now also available for the low-cost Flongle flow cells, which until recently only used the now outdated R9.4 chemistry. This is an important development, because the Flongle is particularly suitable for smaller barcoding projects (200-250 barcodes at a time) that are in high demand.

## Materials and Methods

### PCR with tagged primers

Two sets of amplicons were used to assess the performance of the latest chemistry and real-time barcoding using MinION. The first set consisted of COI amplicons for 7,679 spider specimens while the second consisted of amplicons for 285 phorid specimens also evaluated in Vasilita et al. (2023). DNA was extracted using 20 µl of HotSHOT (total volume = 40 µl after addition of 20 µl neutralisation buffer) (Truett *et al*., 2000). For both experiments tagged amplicons were used. This involved dual tagging where both forward and reverse primers are modified with additional short indices at the 5’ end of the primers. Once amplicons are generated, the reads can be demultiplexed into specimen specific bins after MinION sequencing. Dual tagging allows for multiplexing very large numbers of specimens in single run, i.e. a set of 96 forwards and 96 reverse primers will allow for multiplexing 9,216 products in a single MinION run.

For spiders, a 418-bp fragment of COI was amplified using BF3 (Elbrecht *et al*., 2019) and BR2 (Elbrecht and Leese, 2017) tagged primers. In order to test if short tags can be used 9-bp tags were designed using Barcode Generator (available from http://comailab.genomecenter.ucdavis.edu/index.php/Barcode_generator), with edit-distance of at least 2-bp. PCRs were carried in a reaction mix containing 7 µL of CWBio 2x master mix, 1 µL of BSA, 1 µL of each primer and 3 µL of DNA template, and cycling conditions were as follows: initial denaturation for 15 minutes at 95°C, 25 cycles of: denaturation at 94°C for 30 seconds, annealing 50°C for 1 minutes 30 seconds, extension at 72°C for 1 minutes 30 seconds, and final extension at 72°C for 10 minutes. For phorids, a 313-bp COI fragment was amplified as described in Vasilita et al. (2023). The standard 13-bp tags that we have been routinely using for MinION barcodes as designed for Srivathsan et al. (2019) were used for this set of specimens.

### MinION Sequencing

Amplicons were pooled and prepared for sequencing using MinION. Clean-up was conducted using CleanNGS beads (CleanNA). Library preparation was conducted using Ligation Sequencing Kit V14 (SQK-LSK114, Oxford Nanopore Technologies) with 200 and 100 ng of purified DNA for MinION and Flongle flow cells respectively. The recommended protocol was followed with two modifications as described in Srivathsan et al. (2021). The first modification involves the end-repair reaction which consists of 50□μl of DNA, 7□μl of Ultra II End-prep reaction buffer (New England Biolabs), 3□μl of Ultra II End Prep enzyme mix (New England Biolabs) for a MinION flow cell. The reaction volumes are halved for a Flongle. The second modification was that all bead clean-up steps during library preparation were conducted at a 1X ratio. The amplicon pool corresponding to the spiders was sequenced in a MinION R10.4.1 flow cell and Mk1C at a low-speed setting of 260 bps in order to obtain highly accurate reads with no live basecalling enabled. The amplicon pool corresponding to phorids was sequenced in a Flongle R10.4 and Mk1B with real time barcoding, with live basecalling enabled in MinKNOW. The computer attached to the Mk1B was an 8-core, 32GB RAM workstation with Intel Xeon Processor with RTX 3090 GPU.

### Data Analysis

The overall workflow for MinION based barcoding involves data generation using MinION, basecalling of raw data to obtain a Fastq file with sequences and barcode-calling using ONTbarcoder. Basecalling can be conducted using models and software solutions provided by ONT. There are three main modes of basecalling: “Fast”, “High-accuracy” and “Super-accuracy”, each providing increasingly high per base accuracy at the cost of using more time and GPU-power. For the dataset corresponding to MinION flow cells (spiders), basecalling was conducted using both the fast and super accuracy model using Guppy (Oxford Nanopore Technologies). For Flongle, a second round of basecalling was conducted using high- and super-accuracy models using Guppy after the initial real-time basecalling using the fast model.

In this study, we assessed the performance of real-time barcoding by comparing the resulting barcodes to a reference set of barcodes. This set was obtained using reads called by the super-accuracy model and standard pipeline implemented in ONTbarcoder. This standard pipeline is rigorous in that includes an iterative approach to barcode calling and Quality Controls (QC) that check for barcode length, translatability of the barcode and whether the barcodes contain <1% ambiguous nucleotides. The pipeline is described in detail in Srivathsan et al. (2021) and yields highly accurate barcodes when compared to barcodes obtained with Sanger or Illumina devices. To assess the accuracy of the barcode sets obtained under these various criteria, we used the “Compare barcode sets” module of ONTbarcoder. Lastly to assess the molecular Operational Taxonomic Unit (mOTU) diversity based on the datasets, we used Objective Clustering as implemented in TaxonDNA (Meier *et al*., 2006) using a threshold of 3% (Meier *et al*., 2006; Zhang and Bu, 2022; Srivathsan *et al*., 2023). We used iNEXT to obtain the diversity estimates (Chao et al., 2014; Hsieh and Chao, 2022).

To document the evolution of raw read quality across ONT chemistries (R.9.4, R.10.3, R10.4), the pairwise distance between reads and barcodes was obtained for all demultiplexed reads using the python library *edlib*. The source of data was as follows. The R10.4 reads were obtained here for the amplicon pool consisting of spider sequences (see above). The R10.3 are from Srivathsan et al., 2021: Palaearctic phorid flies; 9,929 specimens and the R9.4 reads from Srivathsan et al., 2019: sample set 1, sequencing run 1, 4,275 specimens. For the latter dataset, the reference barcodes were not constructed using ONTbarcoder because this pipeline is not suitable for ONT’s R9.4 chemistry. Instead, the reference barcodes were obtained with miniBarcoder using extensive error correction (Srivathsan *et al*., 2019). In order to keep the region of comparison consistent across the studies, we trimmed all barcodes to the same 418 bp region of COI obtained for the R10.4 spider data.

Read mapping using *edlib* was done using “HW” or the infix setting to avoid penalizing the gaps at start and end of alignment given that the trimmed consensus barcodes are subsets of the reads.

### Real-time barcoding

We here introduce real-time barcoding as an additional module in ONTbarcoder, available at https://github.com/asrivathsan/ONTbarcoder/releases (see video tutorial: https://youtu.be/nFl8Vw2euMA). In its simplest settings, the new module is lightweight and uses minimal computational resources to avoid interference with an ongoing MinION run, in case the software is used on the same computer as the one used for sequencing. The real-time barcoding module requires two inputs: (1) a demultiplexing file and (2) a link to the folder in which the fastq/fast5 files are saved. The software ensures it analyses any existing data before catching up with sequencing in case ONTbarcoder is not executed in a timely manner.

The configuration for ONTbarcoder depends on whether the user uses a MinION Mk1B or Mk1C for sequencing. If the sequencing is done using Mk1B it is important that the computer connected to the Mk1B can perform basecalling using a GPU. The recommended configurations are described in detail in the section “OPTION 1: Sequencing with Mk1B”. If sequencing involves a Mk1C, one can either use the internal or an external GPU server for basecalling. However, the built-in GPU currently can only keep up with basecalling when using the fast model. More details can be found in the section on “OPTION 2: Sequencing with Mk1C”. An alternative is to pair a Mk1C with a computer that has a GPU that is powerful enough to use higher accuracy basecalling models and yet keeps up with data generation. This is described under “Mk1C with external basecalling”. Overall, we recommend the use of Mk1C if one is available because it allows for controlling the temperature, which is needed for the 260 bps setting of R10.4 flow cells. If only a Mk1B is available, we recommend using the 400 bps setting.

#### OPTION 1: Sequencing with Mk1B

Real-time barcoding using Mk1B requires connecting a computer that can perform GPU-based basecalling. MinKNOW and ONTbarcoder should be installed on the same computer. If the computer has a slow GPU, only the fast basecalling model may keep up with data generation. More powerful GPUs allow for the use of high-or super-accuracy models. Basecalling can either be conducted live via MinKNOW or via ONTbarcoder. If the latter is chosen, ONTbarcoder will configure basecalling via Guppy and hence Guppy (gpu-version) must be installed on the computer.

#### OPTION 2: Sequencing with Mk1C

An external GPU is not essential for this option (shown in video tutorial: https://youtu.be/nFl8Vw2euMA). Instead, a standard computer can analyze the data generated by the Mk1C and then transferred to the computer running ONTbarcoder. In this case, MinKNOW in Mk1C configures and manages the data generation and basecalling while ONTbarcoder is installed on a separate computer and connects over the network (preferably ethernet or hotspot). ONTbarcoder will monitor and transfer fastq from the Mk1C and then carry out the real-time demultiplexing and barcode calling.

Mk1C with external basecalling: This is an advanced configuration that is recommended for a user who wants to take advantage of a computer with a powerful GPU. Such a computer allows for real-time high-or super-accuracy basecalling. In this configuration, ONTbarcoder connects to the remote device Mk1C, transfers the fast5 files as they are generated. GPU-enabled basecalling is done using the computer that also hosts ONTbarcoder. ONTbarcoder allows the user to configure Guppy for basecalling and manages four real time processes; i.e., file transfers, basecalling (via Guppy), demultiplexing, and barcode-calling.

To enable these, different real-time processes are supported in ONTbarcoder. The two main parallel processes are demultiplexing and barcode calling. One thread monitors the generation of fastq files. Identification of a new file pauses the monitoring until all reads in the new file are demultiplexed into specimen-specific bins. By default, ONT stores 1,000 (Flongle) or 4,000 (MinION) reads per file. Once a file is demultiplexed, the file tracking for new fastq files resumes. A second thread of the real-time barcoding module monitors how many reads are already available for a particular specimen. This count is updated after the demultiplexing of a file is completed. All specimen bins exceeding a minimum coverage (default=20) are identified and barcodes are called. All barcodes that satisfy the QC-criteria are accepted and removed from the list of specimens that need barcodes. All barcodes that do not satisfy ONTbarcoder’s QC criteria are rejected and barcode calling is tried again after additional reads have been gathered. Once a round of consensus calling is completed the thread monitoring coverage resumes, and identifies the new specimens that satisfy minimum coverage, or those specimens that failed QC but gathered additional reads.

The real-time barcoding module uses the following additional feature to avoid slowing down barcode calling. Firstly, specimen bins that fail to yield barcodes at medium coverage (e.g., 50 reads) often also do not yield MinION barcodes at very high coverage due to unclear signal in the data (Srivathsan *et al*., 2021). However, revisiting such reads sets every time after new reads have been demultiplexed is computationally expensive. Therefore, all reads sets that failed to yield a barcode at high coverage (default=100) are only revisited when multiple new reads are available. The default step size is 5 so that the barcode would only be called again after 5 additional reads are accumulated unless the defaults are modified. Secondly, the real-time barcoding module runs faster when a maximum read coverage is specified, because bins with very high coverage slow down the alignment step. This feature is turned off by default, but if activated only a user-specified number of reads are used for barcode calling. For this purpose, the reads closest to the expected length of the barcode are selected.

The other two real-time features of ONTbarcoder are real-time file transfers and real-time basecalling via guppy. These are implemented in a manner similar to demultiplexing and barcode calling where a thread monitors progress, pauses while the task is conducted and resumes again. The real-time file transfer module monitors the fastq/fast5 file generation in a remote folder and transfers the files over as they are generated. So far, we have observed at least a 20 second gap between generation of different files (1000-4000 sequences in a fast5/fastq). Thus, this module tends to keep up with the data generation even over hotspot connections.

The live basecalling module configures Guppy executable of the system and passes the fast5 files to be basecalled. Within this procedure, a batch of fast5 files available at the time of completion of a previous round of basecalling are basecalled together.

For demultiplexing, ONTbarcoder constructs a library of tags and tag variants upon reading the demultiplexing file prior to the start of real time barcoding. For consensus barcode calling, ONTbarcoder uses MAFFT v7 (Katoh and Standley, 2013) with parameters trained for MinION reads to construct a Multiple Sequence Alignment (Hamada *et al*., 2017; Frith *et al*., 2021). A barcode is called using an iterative consensus criterion starting at a minimum frequency of 0.3 for each position (empirically determined in Srivathsan et al. 2021). If using the default does not yield a barcode that satisfies the QC criteria, the frequency is varied from 0.2 to 0.5.

The results for the real time barcoding are saved in a folder which contains a subfolder for demultiplexed reads. The output consensus sequences are stored in the “consensus_good.fa” which is updated after every iteration of consensus calling. For specimens that only yielded barcodes that failed the QC, the result of the last attempt to call the barcode is saved in the “consensus_all.fa” which is routinely updated to keep the latest consensus barcodes. This file includes the sequences that passed QC controls.

## Results

### Performance of R10.4 chemistry for barcoding

We sequenced 7,679 specimens and 81 negatives using an R10.4 flowcell and obtained 14,489,886 sequences, of which 9,448,704 were in the 50-bp window of the expected product length. These sequences were first basecalled using the super-accuracy model. Demultiplexing and barcode calling using the standard pipeline (“Conventional pipeline”) of ONTbarcoder yielded 4,230,457 (44.8%) demultiplexed reads from which 6,950 barcodes could be called, giving an overall barcoding success rate of 90.5%. This is higher than the PCR success estimated by agarose gel electrophoresis for a subset of samples (81.8%). Thus, the new ONT chemistry allows for the use of shorter tags and reducing the error tolerance to 1-bp.

We next examined the quality of reads basecalled under the super accuracy (sup) model by mapping the reads back onto the consensus barcodes. R10.4 reads are very accurate with 25 % of the reads having just <=1 bp error while 54% of the reads have fewer than 4 bp errors or >99% accuracy (N50>99%) (Figure 1). In comparison, the N50 for R10.3 (HAC) was 94% and R9.4 (Fast) was 89.2%. Improvements in quality also resulted in a higher proportion of the reads being demultiplexed (Figure 2).

**Figure 1:**
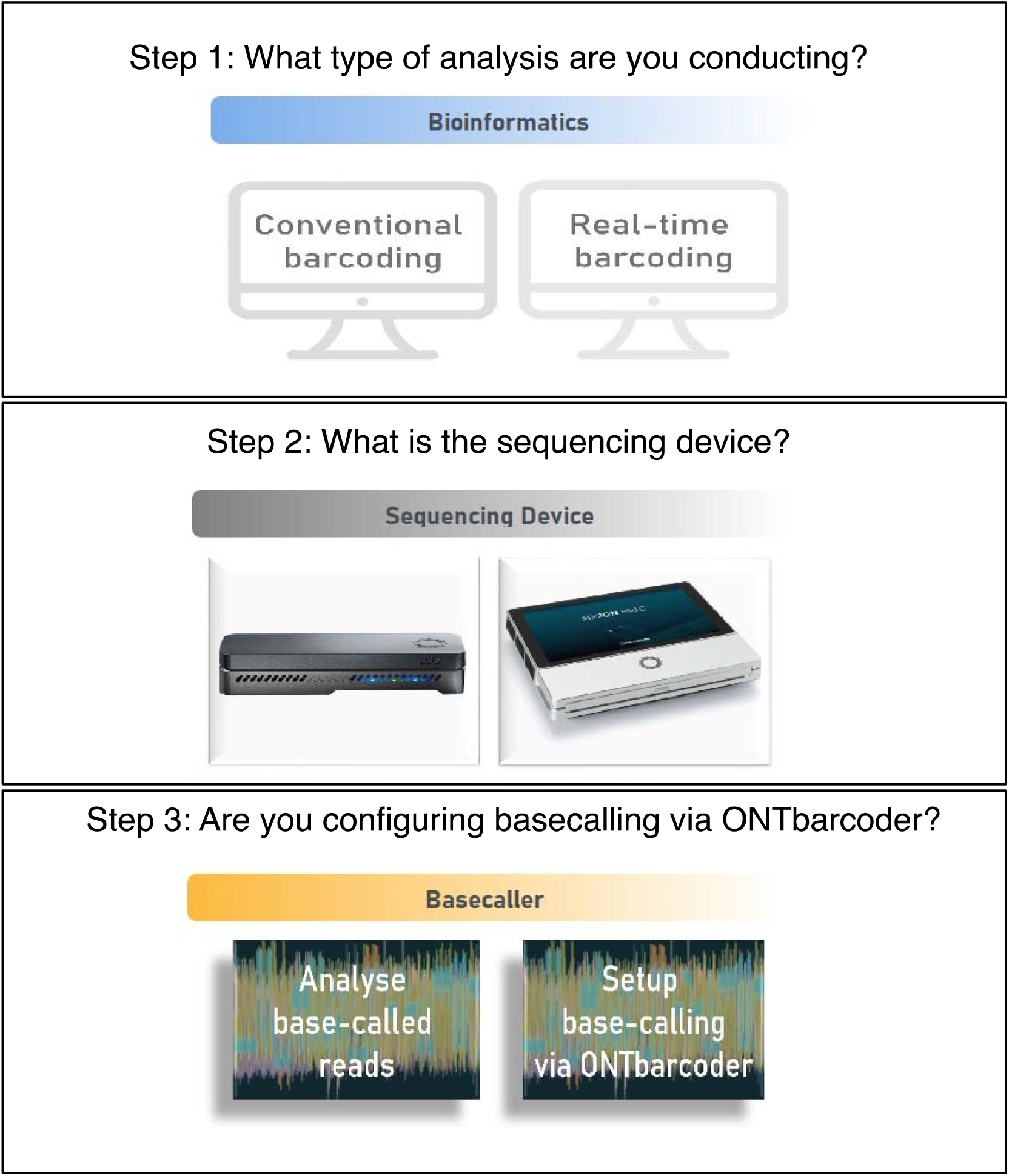
Options for the initial configuration via ONTbarcoder

**Figure 2:**
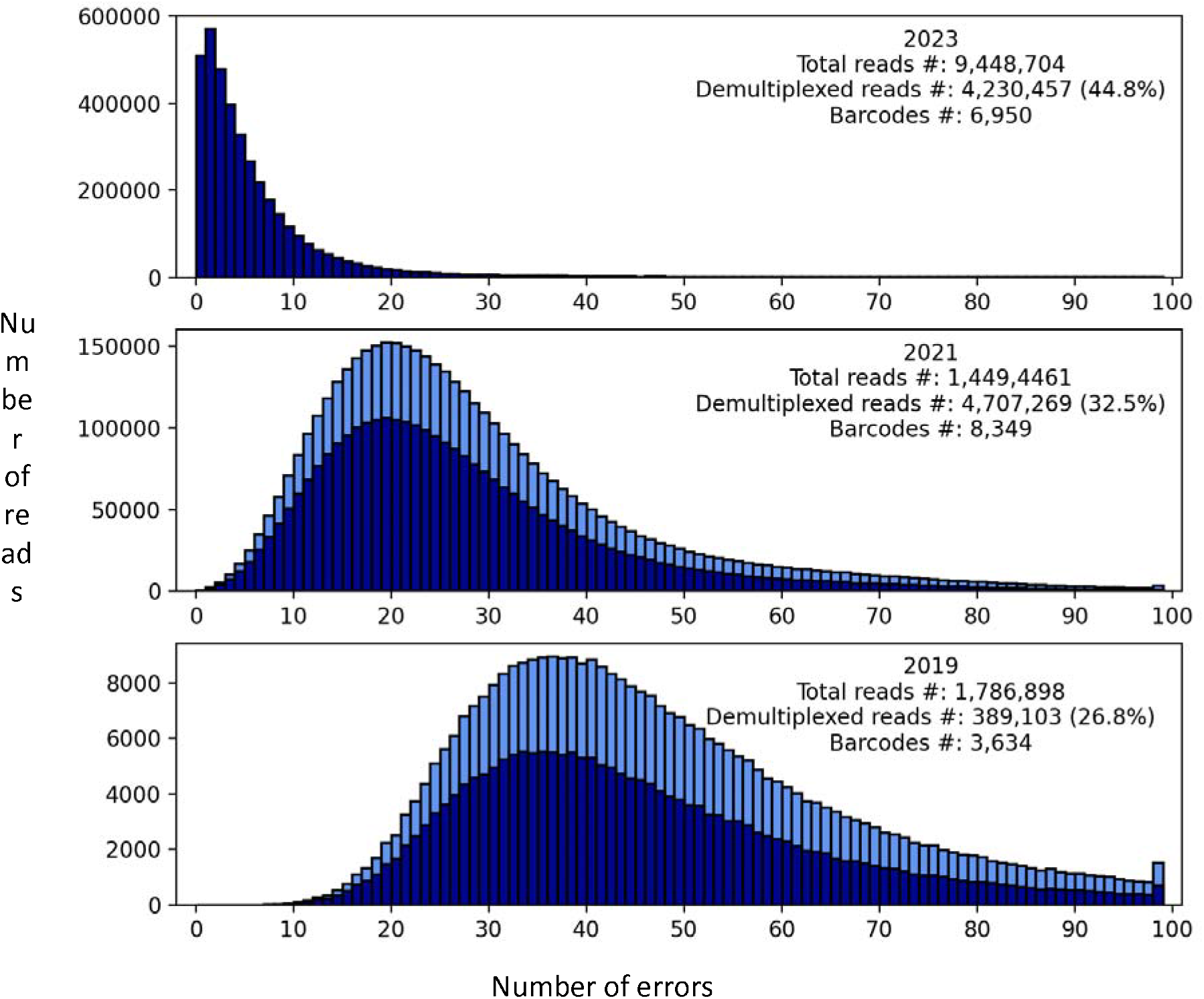
Number of errors per raw read when reads are mapped to consensus barcodes. The dark and light blue represents the reads demultiplexed when allowing for a 1-bp or 2-bp error in the primer tag, respectively. Demultiplexing for R10.4 data in 2023 was conducted at only 1-bp setting.

The improvement in raw read quality also enhanced barcoding accuracy. Firstly, the proportion of specimen bins with >5X coverage yielding high quality barcodes is much higher for R10.4 (94.5% for R10.4+sup, 88.6% for R10.3+hac and 72.4% for R9.4+fast). Thus, short tags do not lead to an increased amount of incorrectly demultiplexed reads and overall, there is an improvement in the accuracy of specimen binning. Secondly, more barcodes are obtained without using all iterative error correction tools that are part of the the original version of ONTbarcoder (Table 1). With R10.4 chemistry and using the super-accuracy basecalling model, 97.2% of the barcodes are obtained in the first step of consensus calling and only 0.3% of the barcodes require any indel correction.

**Table 1:** Number of barcodes obtained at various stages of ONTbarcoder’s pipeline based on the latest chemistry and basecalling model available for analysis at the time of sequencing.

| Dataset | Consensus by length | Consensus by similarity | Consensus by barcode comparison | # Final filtered barcodes |
| --- | --- | --- | --- | --- |
| R10.4+sup | 6,754 (97.2%) | 178 (2.6%) | 18 (0.3%) | 6,950 |
| R10.3+hac | 7,630 (91.4%) | 341 (4.1%) | 377 (4.5%) | 8,348 |
| R9.4+fast | 288 (11.4%) | 119 (4.7%) | 2,112 (83.8%) | 2,519 |

Given the results obtained with the super-accuracy model, we next examined if fast-basecalling R10.4 reads could also yield good quality barcodes. We found 6,650 good barcodes using ONTbarcoder’s default pipeline, giving a barcoding success rate of 86.6%. This is lower than what was obtained with super-accuracy basecalling (90.5%), but higher than the estimated PCR success rate based on electrophoresis gels. A pairwise comparison between barcodes obtained using fast and super-accuracy models showed an overall similarity of 99.93%. Thus, reads called using fast-basecalling are able to yield accurate barcodes.

### Real-time barcoding with Flongle

We sequenced a pool of 285 phorid fly PCR products using a Flongle with R10.4 chemistry using a MinION Mk1B attached to a GPU powered computer. Real-time barcoding using reads obtained with fast-basecalling yielded 172 barcodes (Figure 3). More than 100 barcodes were produced within the first two hours of sequencing with little further gain after four hours. Note that the original live barcoding run was conducted with an earlier version of the GUI (see results in Supplementary Figure 1). Figure 1 is reconstructed based on an updated GUI and simulation based on time of data generation.

**Figure 3:**
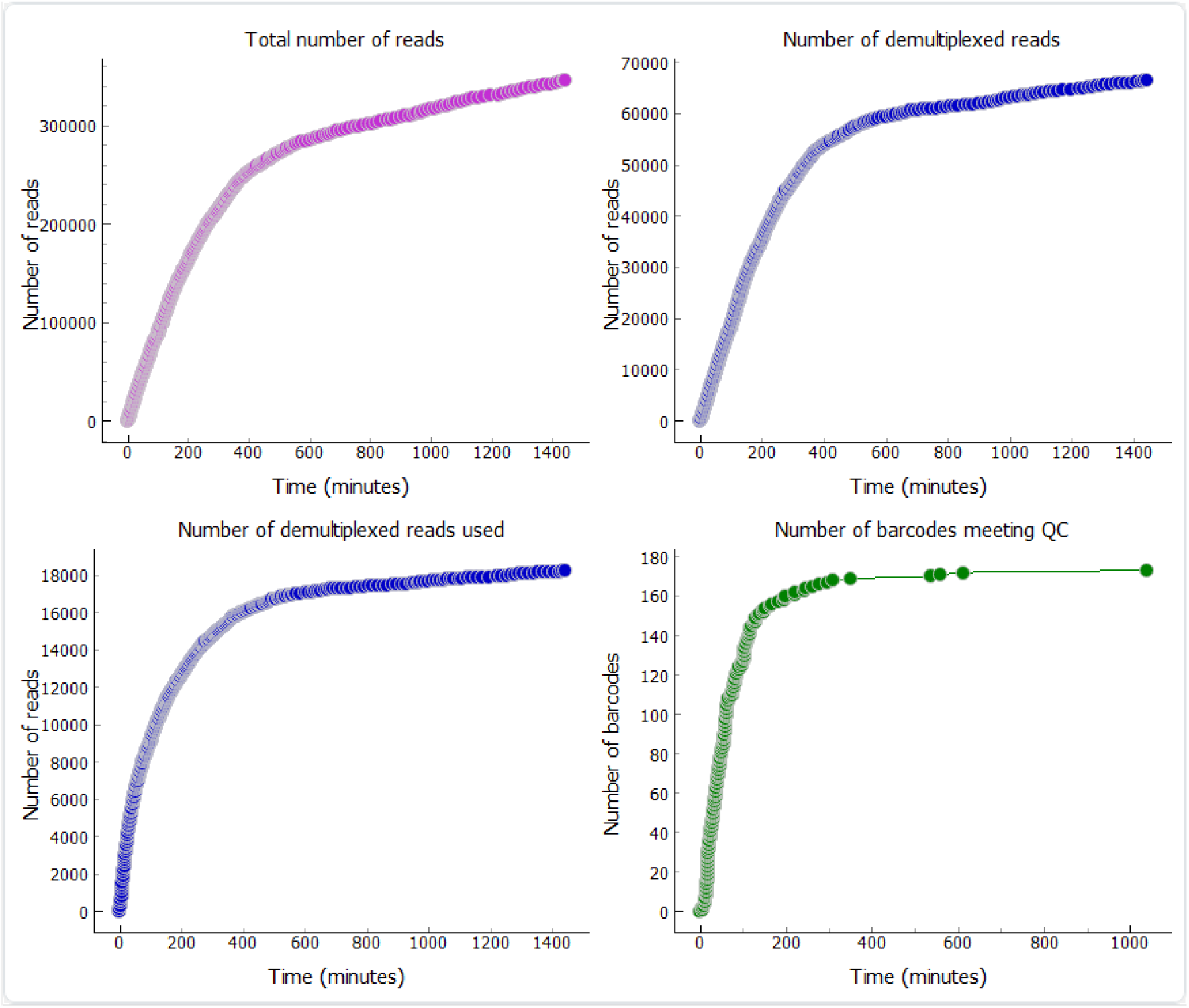
A sample real time barcoding run using the Flongle R10.4 flow cell in ONTbarcoder using fast basecalling. The four panels in the software keep track of the following: top left-total number of basecalled reads, top right-total number of reads demultiplexed, bottom left-number of demultiplexed reads used for barcode calling, and; bottom right: number of barcodes called. The x-axis in all four graphs is time in minutes.

We subsequently re-analyzed the same data using high- and super-accuracy basecalling models by simulating real-time barcoding based on time-stamps. The use of the higher quality basecallers increased the number of barcodes from 172 (71%) to 220 (91%) and 224 (93%) barcodes respectively (Table 2, Supplementary Figures 2&3). The higher accuracy basecalling models also improved the quality of the consensus barcodes. Pairwise comparison of the barcodes obtained with real-time barcoding with the final reference barcodes revealed a per base accuracy of 99.7%, 99.98% and 99.98% for the barcodes based on reads called by fast, high- and super-accuracy basecalling. Lastly, these models yielded QC-compliant barcodes faster. For results corresponding to both the high accuracy and super-accuracy models, >180 barcodes could be obtained within two hours of sequencing.

**Table 2:**
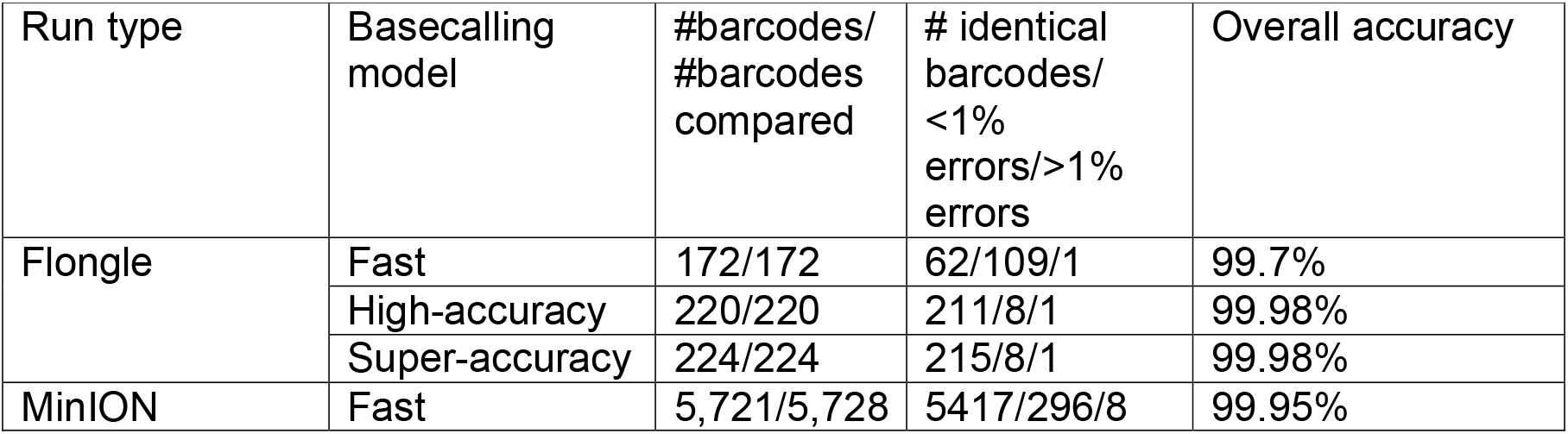
Accuracy of barcodes obtained using real-time barcoding module.

| Run type | Basecalling model | #barcodes/<br>#barcodes compared | # identical barcodes/<br><1% errors/>1% errors | Overall accuracy |
| --- | --- | --- | --- | --- |
| Flongle | Fast | 172/172 | 62/109/1 | 99.7% |
|  | High-accuracy | 220/220 | 211/8/1 | 99.98% |
|  | Super-accuracy | 224/224 | 215/8/1 | 99.98% |
| MinION | Fast | 5,721/5,728 | 5417/296/8 | 99.95% |

We next assessed how the different sets of barcodes influenced the assessment of putative species diversity in a sample. We found that real-time barcoding based on fast basecalling yielded 22 mOTUs, high-accuracy reads yielded 25 mOTUs and super-accuracy reads yielded 25 mOTUs. In comparison the standard pipeline yielded 29 mOTUs. With regard to species accumulation curves, the diversity estimates have overlapping confidence intervals (Figure 4) suggesting that a preliminary diversity profile can be obtained using real-time barcoding regardless of which basecaller is used. However, some species were only detected in the most comprehensive analysis of all data.

**Figure 4:**
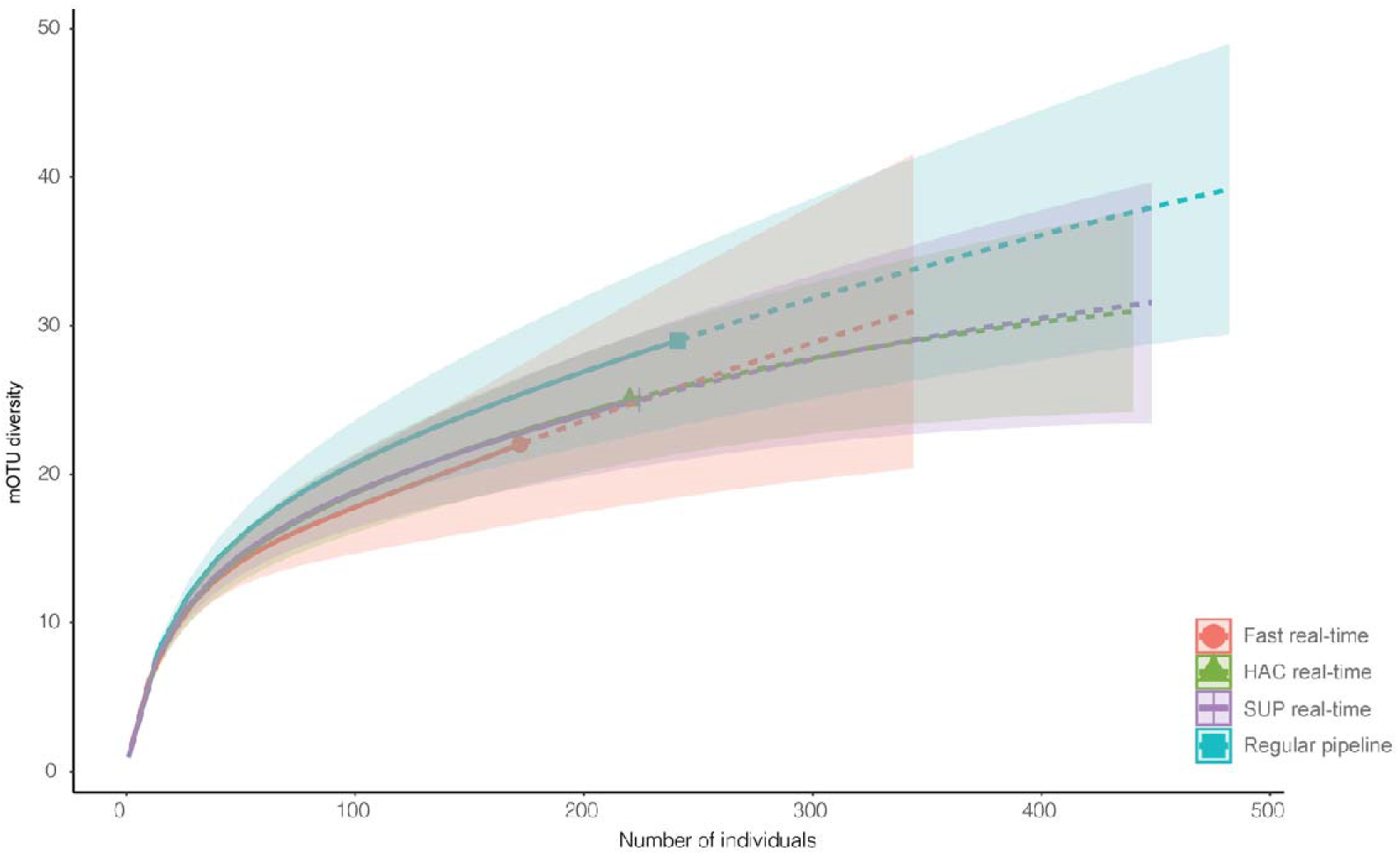
mOTU accumulation curves showing overlapping diversity estimates of different datasets obtained from the sample of phorids sequenced using a single Flongle run. The solid lines represents interpolation while dotted lines represents extrapolation.

### Real-time barcoding with MinION

We assessed if real-time barcoding with fast-basedcalled reads would yield accurate results for the full MinION run by using the time stamps of the R10.4 run. A total 5,728 barcodes could be called using real time barcoding i.e. 82% of the final barcodes obtained using super-accuracy model and standard ONTbarcoder pipeline (Figure 5). The accuracy of the barcodes is 99.95% when comparing them with the 6,950 barcodes obtained using the standard pipeline of ONTbarcoder applied to the reads obtained with super-accuracy basecalling. Real time barcoding yielded 50% of the barcodes within 3 hours and 80% within 10 hours of sequencing. For both Flongle and MinION, the number of reads used for barcoding and number of barcodes accumulated to a plateau much faster than the total number of reads and demultiplexed reads; i.e., the reads obtained later in the run are of little use for constructing consensus barcodes because many reads pertain to specimens whose barcode was called earlier. In terms of mOTU diversity, at 3%, we found that the standard pipeline yielded 197 mOTUs whereas real-time barcoding based on fast basecalled reads yielded 179 mOTUs.

**Figure 5:**
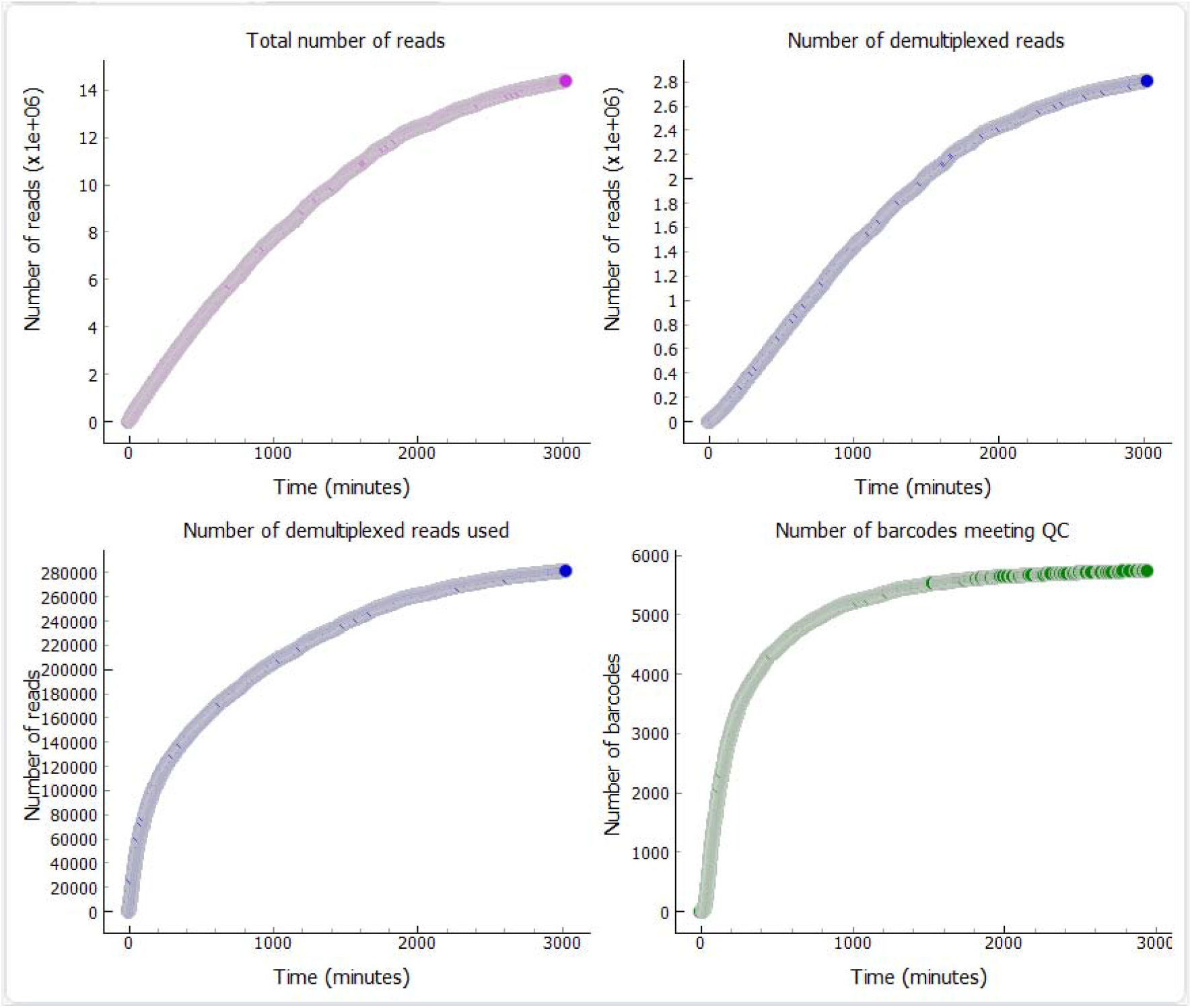
Real time barcoding using MinION R10.4 flowcell for 7679 spiders. More than 5,700 barcodes were obtained with bulk of the barcodes called within 10 hours of sequencing.

## Discussion

Species discovery and identification should ideally be simple, cost-effective, fast, and accurate. DNA barcoding with ONT’s MinION gets us closer to this goal because it is rapid and cost-effective (Pomerantz *et al*., 2018; Srivathsan *et al*., 2018; Maestri *et al*., 2019). However, until now there was no bioinformatics pipeline that was able to generate real-time species richness estimates and identifications to match the real-time read generation. Moreover, many bioinformatics tools were not user-friendly, which will interfere with making MinION barcoding popular. In the past, real-time barcoding was difficult to implement because of the high error rates of MinION sequencers in the past (Figure 2). Here, we demonstrate that the improvements in the quality of MinION data are such that real-time barcoding is no longer elusive.

We show that real-time barcoding with reads obtained using fast basecalling yields 70-80% of the final barcodes and 75-90% of the final mOTU diversity as long as R10.4 reads are used. Thus, one can obtain a good species diversity profile based on a preliminary analysis of the reads while sequencing is ongoing (Figure 4). All common species are discovered within minutes and only rare species represented by few reads are only added when analyzing all reads obtained with the best basecallers. If the user has access to a computer with GPUs that allow for real-time basecalling using high-or super-accuracy models, the results can be even better allowing for the recovery of >90% of individual barcodes and >85% of the mOTU diversity. Furthermore, the use of better base-calling models also improve barcode quality (99.98%), although the improvement is moderate given that the accuracy of barcodes obtained with fast basecalling is already ∼99.9%. Note, however, that we nevertheless recommend that the results based on real-time barcoding are only considered preliminary because an analysis of all data allows for applying the best base-calling, selecting reads from a larger pool, and the application of several iterative correction steps. This means that the post-hoc analysis of all data is more likely to recover barcodes for very weak amplicons.

To further speed up the process of estimating the species diversity in a bulk sample, real-time barcoding can be coupled with the “express barcoding” of Vasilita et al. (2023) that accelerates amplicon production by using NextGenPCR® thermocyclers. They use special polymerases and fast heating/cooling in three preset heating zones. Thus, the PCR amplification of a 313-bp fragment of COI in 30 cycles only requires 22 minutes. With a single such machine, Vasilita et al. (2023) went from specimens to identified barcodes for 285 phorid specimens in 6 hours. With real-time barcoding, a preliminary analysis of the barcodes can be conducted as the reads are generated. Barcodes can be called and identified using SequenceID (https://www.gbif.org/tools/sequence-id) within minutes of starting a MinION run.

A particularly useful feature of real-time data analysis is that it allows the users to monitor the progress of large-scale barcoding experiments and make decisions regarding when to stop sequencing. For instance, we show that barcode accumulation slows dramatically after ca. three hours of sequencing on a Flongle. A re-analysis of the dataset using super-accuracy model for reads accumulated during the first three hours shows that the overall success rate only drops by 2% success compared to analyzing all reads for a 24-hour run (Vasilita *et al*., 2023, Table 1).

Thus, users can make informed choices on which coverage is needed for any particular project. This is particularly useful for studies where only the dominant species in a specimen sample needs to be identified. Stopping a sequencing run after a few hours, allows for re-using the same flow cell for future projects.

Real-time barcoding presents itself as a useful platform to encourage citizen scientist and pre-tertiary student involvement in species discovery projects. DNA barcodes allow for involving students directly in the process of species discovery in a systematic manner (Chiovitti *et al*., 2019). However, this requires that barcoding is sufficiently fast and requires limited equipment (Marizzi *et al*., 2018). Real-time barcoding is here useful because it delivers barcodes fast and requires little equipment. Indeed, real-time barcoding could also be an important element of science or natural history museum exhibition on species discovery. It could be shown how specimens of a bulk sample go through three stages. Robotic imaging with a robot such as the DiversityScanner (Wührl et al., 2021) followed by real-time barcoding and real-time species identification with a software tool such as GBIF’s SequenceID (https://www.gbif.org/tools/sequence-id).

Other applications of rapid barcoding include the identification of unknown species in the field (Pomerantz *et al*., 2018; Maestri *et al*., 2019), detection of illegal trade (Seah *et al*., 2020), detection of insect outbreaks (Cranston *et al*., 2013), food authentication (Ho *et al*., 2020) and non-human forensics (Vasiljevic *et al*., 2021).

The new features in ONTbarcoder were made feasible due to the rapid improvement in ONT read quality which in turn improved demultiplexing rates and reduced the number of barcodes that required any kind of error correction. It is noteworthy that the developments in chemistry are not limited to MinION. The availability of R10 technology for Flongle makes the chemistry accessible for users interested in sequencing smaller number of specimens. Moreover, the improvements allowed for reducing primer tag length of the primers, which should improve PCR efficiency. Note that ONTbarcoder 2.0 now allows for modifying the number of permissible errors in the tags.

### Conclusions

We here document how MinION generated data have improved over time.

Looking ahead, such improvements are likely to continue. In addition to the developments discussed in this study, a major step forward has come in the form of the P2 sequencing device. Each PromethION flow-cell contains 5X number of nanopore channels compared to a MinION flow-cell even though the cost of the two types of flow-cells is similar. Overall, these developments make it likely that ONT barcoding will have a bright future and can boost biodiversity discovery and biomonitoring when it is needed most.

## Supporting information

Supplementary Table Primer Tags

## Acknowledgements

We would like to thank Cristina Vasilita for the initial work on the set of phorid specimens that were sequenced during the express barcoding project. We would also like to thank Mary Ann Davenport, Shauna Kehoe and Denise Cheah for their help with library preparation. We thank Michał Kolasa and Piotr Łukasik for the PCR reaction conditions. Daniel Suárez was funded by the ‘Ministerio de Ciencia e Innovación’ through an FPI PhD fellowship (PRE2018-083230). Spider sampling was supported by the project CGL2017-85718-P (funded by MCIN/AEI/ 10.13039/501100011033, Spain and EDRF, EU) awarded to Brent C. Emerson.

Spider fieldwork was conducted with the collecting permits 246285, AFF-107/17 (sigma no. 2017-00572), A/EST-034/16, RE:2349 and REUS-11227 kindly provided by the Canary Islands Government, Cabildo of Tenerife, Cabildo of La Palma, Cabildo of El Hierro, and Cabildo de La Gomera, respectively.

## Author Contributions

RM and AS conceptualized the study, AS developed the software, VF and DS performed molecular work, VF tested the software, DS and BE collected the samples. AS and RM wrote the manuscript and all authors read and approved the manuscript.

## Data Availability Statement

ONTbarcoder2 is available https://github.com/asrivathsan/ONTbarcoder/releases. The data have been uploaded to Figshare and will be available at 10.6084/m9.figshare.23826378. The private link for review is: https://figshare.com/s/53b0b0ce6978f658146c.

## References

Bar-On, Y.M., Phillips, R., Milo, R. 2018. The biomass distribution on Earth. Proceedings of the National Academy of Sciences. 115, 6506–6511. doi:10.1073/pnas.1711842115

Belovsky, G.E., Slade, J.B. 2000. Insect herbivory accelerates nutrient cycling and increases plant production. Proceedings of the National Academy of Sciences. 97, 14412–14417. doi:10.1073/pnas.250483797

Bianchi, F.J.J.A., Booij, C.J.H., Tscharntke, T. 2006. Sustainable pest regulation in agricultural landscapes: a review on landscape composition, biodiversity and natural pest control. Proceedings of the Royal Society B: Biological Sciences. 273, 1715–1727. doi:10.1098/rspb.2006.3530

Biesmeijer, J.C., Roberts, S.P.M., Reemer, M., Ohlemulller, R., Edwards, M., Peeters, T., Schaffers, A.P., Potts, S.G., Kleukers, R., Thomas, C.D., Settele, J., Kunin, W.E. 2006. Parallel declines in pollinators and insect-pollinated plants in Britain and the Netherlands. Science. 313, 351–354. doi:10.1126/science.1127863

Bohmann, K., Elbrecht, V., Carøe, C., Bista, I., Leese, F., Bunce, M., Yu, D.W., Seymour, M., Dumbrell, A.J., Creer, S. 2022. Strategies for sample labelling and library preparation in DNA metabarcoding studies. Molecular Ecology Resources. 22, 1231–1246. doi:10.1111/1755-0998.13512

Chao, A., Gotelli, N.J., Hsieh, T.C., Sande, E.L., Ma, K.H., Colwell, R.K., Ellison, A.M. 2014. Rarefaction and extrapolation with Hill numbers: a framework for sampling and estimation in species diversity studies.. Ecological Monographs. 84, 45–67.

Chiovitti, A., Thorpe, F., Gorman, C., Cuxson, J.L., Robevska, G., Szwed, C., Duncan, J.C., Vanyai, H.K., Cross, J., Siemering, K.R., Sumner, J. 2019. A citizen science model for implementing statewide educational DNA barcoding. PLOS ONE. 14, e0208604. doi:10.1371/journal.pone.0208604

Chua, P.Y.S., Bourlat, S.J., Ferguson, C., Korlevic, P., Zhao, L., Ekrem, T., Meier, R., Lawniczak, M.K.N. 2023. Future of DNA-based insect monitoring. Trends in Genetics. doi:10.1016/j.tig.2023.02.012

Cranston, P.S., Ang, Y.C., Heyzer, A., Lim, R.B.H., Wong, W.H., Woodford, J.M., Meier, R. 2013. The nuisance midges (Diptera: Chironomidae) of Singapore’s Pandan and Bedok Reservoirs. The Raffles Bulletin of Zoology. 61, 779–793.

Elbrecht, V., Braukmann, T.W.A., Ivanova, N. V., Prosser, S.W.J., Hajibabaei, M., Wright, M., Zakharov, E. V., Hebert, P.D.N., Steinke, D. 2019. Validation of COI metabarcoding primers for terrestrial arthropods. PeerJ. 7, e7745. doi:10.7717/peerj.7745

Elbrecht, V., Leese, F. 2017. Validation and development of COI metabarcoding primers for freshwater Macroinvertebrate Bioassessment. Frontiers in Environmental Science. 5. doi:10.3389/fenvs.2017.00011

Frith, M.C., Mitsuhashi, S., Katoh, K. 2021. lamassemble: Multiple Alignment and Consensus Sequence of Long Reads. pp. 135–145. doi:10.1007/978-1-0716-1036-7_9

Fürst, J., Bollmann, K., Gossner, M.M., Duelli, P., Obrist, M.K. 2023. Increased arthropod biomass, abundance and species richness in an agricultural landscape after 32 years. Journal of Insect Conservation. 27, 219–232. doi:10.1007/s10841-022-00445-9

Gallai, N., Salles, J.-M., Settele, J., Vaissière, B.E. 2009. Economic valuation of the vulnerability of world agriculture confronted with pollinator decline. Ecological Economics. 68, 810–821. doi:10.1016/j.ecolecon.2008.06.014

Hamada, M., Ono, Y., Asai, K., Frith, M.C. 2017. Training alignment parameters for arbitrary sequencers with LAST-TRAIN. Bioinformatics. 33, 926–928. doi:10.1093/bioinformatics/btw742

Hebert, P.D.N., Cywinska, A., Ball, S.L., deWaard, J.R. 2003. Biological identifications through DNA barcodes. Proceedings of the Royal Society of London. Series B: Biological Sciences. 270, 313–321. doi:10.1098/rspb.2002.2218

Ho, J.K.I., Puniamoorthy, J., Srivathsan, A., Meier, R. 2020. MinION sequencing of seafood in Singapore reveals creatively labelled flatfishes, confused roe, pig DNA in squid balls, and phantom crustaceans. Food Control. 112, 107144. doi:10.1016/j.foodcont.2020.107144

Hsieh, T.C., Chao, A. 2022. iNEXT: Interpolation and Extrapolation for Species Diversity.

Katoh, K., Standley, D.M. 2013. MAFFT Multiple Sequence Alignment Software version 7: Improvements in performance and usability. Molecular Biology and Evolution. 30, 772–780. doi:10.1093/molbev/mst010

Maestri, Cosentino, Paterno, Freitag, Garces, Marcolungo, Alfano, Njunjić, Schilthuizen, Slik, Menegon, Rossato, Delledonne. 2019. A rapid and accurate MinION-based workflow for tracking species biodiversity in the field. Genes. 10, 468. doi:10.3390/genes10060468

Marizzi, C., Florio, A., Lee, M., Khalfan, M., Ghiban, C., Nash, B., Dorey, J., McKenzie, S., Mazza, C., Cellini, F., Baria, C., Bepat, R., Cosentino, L., Dvorak, A., Gacevic, A., Guzman-Moumtzis, C., Heller, F., Holt, N.A., Horenstein, J., Joralemon, V., Kaur, M., Kaur, T., Khan, A., Kuppan, J., Laverty, S., Lock, C., Pena, M., Petrychyn, I., Puthenkalam, I., Ram, D., Ramos, A., Scoca, N., Sin, R., Gonzalez, I., Thakur, A., Usmanov, H., Han, K., Wu, A., Zhu, T., Micklos, D.A. 2018. DNA barcoding Brooklyn (New York): A first assessment of biodiversity in Marine Park by citizen scientists. PLOS ONE. 13, e0199015. doi:10.1371/journal.pone.0199015

Meier, R., Shiyang, K., Vaidya, G., Ng, P.K.L. 2006. DNA barcoding and taxonomy in Diptera: A tale of high intraspecific variability and low identification success. Systematic Biology. 55, 715–728. doi:10.1080/10635150600969864

Meier, R., Wong, W., Srivathsan, A., Foo, M. 2016. $1 DNA barcodes for reconstructing complex phenomes and finding rare species in specimen-rich samples. Cladistics. 32, 100–110. doi:10.1111/cla.12115

Oliver, T.H., Isaac, N.J.B., August, T.A., Woodcock, B.A., Roy, D.B., Bullock, J.M. 2015. Declining resilience of ecosystem functions under biodiversity loss. Nature Communications. 6, 10122. doi:10.1038/ncomms10122

Outhwaite, C.L., McCann, P., Newbold, T. 2022. Agriculture and climate change are reshaping insect biodiversity worldwide. Nature. 605, 97–102. doi:10.1038/s41586-022-04644-x

Pomerantz, A., Peñafiel, N., Arteaga, A., Bustamante, L., Pichardo, F., Coloma, L.A., Barrio-Amorós, C.L., Salazar-Valenzuela, D., Prost, S. 2018. Real-time DNA barcoding in a rainforest using nanopore sequencing: opportunities for rapid biodiversity assessments and local capacity building. GigaScience. 7. doi:10.1093/gigascience/giy033

Pomerantz, A., Sahlin, K., Vasiljevic, N., Seah, A., Lim, M., Humble, E., Kennedy, S., Krehenwinkel, H., Winter, S., Ogden, R., Prost, S. 2022. Rapid in situ identification of biological specimens via DNA amplicon sequencing using miniaturized laboratory equipment. Nature Protocols. 17, 1415–1443. doi:10.1038/s41596-022-00682-x

Ratnasingham, S., Hebert, P.D.N. 2013. A DNA-based registry for all animal species: The Barcode Index Number (BIN) system. PLoS ONE. 8, e66213. doi:10.1371/journal.pone.0066213

Schmeller, D.S., Courchamp, F., Killeen, G. 2020. Biodiversity loss, emerging pathogens and human health risks. Biodiversity and Conservation. 29, 3095–3102. doi:10.1007/s10531-020-02021-6

Seah, A., Lim, M.C.W., McAloose, D., Prost, S., Seimon, T.A. 2020. MinION-Based DNA barcoding of preserved and non-invasively collected wildlife samples. Genes. 11, 445. doi:10.3390/genes11040445

Srivathsan, A., Ang, Y., Heraty, J.M., Hwang, W.S., Jusoh, W.F.A., Kutty, S.N., Puniamoorthy, J., Yeo, D., Roslin, T., Meier, R. 2023. Convergence of dominance and neglect in flying insect diversity. Nature Ecology & Evolution. doi:10.1038/s41559-023-02066-0

Srivathsan, A., Baloğlu, B., Wang, W., Tan, W.X., Bertrand, D., Ng, A.H.Q., Boey, E.J.H., Koh, J.J.Y., Nagarajan, N., Meier, R. 2018. A MinION^TM^-based pipeline for fast and cost-effective DNA barcoding. Molecular Ecology Resources. 18, 1035–1049. doi:10.1111/1755-0998.12890

Srivathsan, A., Hartop, E., Puniamoorthy, J., Lee, W.T., Kutty, S.N., Kurina, O., Meier, R. 2019. Rapid, large-scale species discovery in hyperdiverse taxa using 1D MinION sequencing. BMC Biology. 17. doi:10.1186/s12915-019-0706-9

Srivathsan, A., Lee, L., Katoh, K., Hartop, E., Kutty, S.N., Wong, J., Yeo, D., Meier, R. 2021. ONTbarcoder and MinION barcodes aid biodiversity discovery and identification by everyone, for everyone. BMC Biology. 19. doi:10.1186/s12915-021-01141-x

Tallamy, D.W., Shriver, W.G. 2021. Are declines in insects and insectivorous birds related?. Ornithological Applications. 123. doi:10.1093/ornithapp/duaa059

Truett, G.E., Heeger, P., Mynatt, R.L., Truett, A.A., Walker, J.A., Warman, M.L. 2000. Preparation of PCR-quality mouse genomic DNA with hot Sodium Hydroxide and Tris (HotSHOT). BioTechniques. 29, 52–54. doi:10.2144/00291bm09

Vasilita, C., Feng, V., Hansen, A.K., Hartop, E., Srivathsan, A., Struijk, R., Meier, R. Express barcoding with NextGenPCR and MinION for species-level sorting of ecological samples. bioRxiv. doi:10.1101/2023.04.27.538648

Vasiljevic, N., Lim, M., Humble, E., Seah, A., Kratzer, A., Morf, N. V., Prost, S., Ogden, R. 2021. Developmental validation of Oxford Nanopore Technology MinION sequence data and the NGSpeciesID bioinformatic pipeline for forensic genetic species identification. Forensic Science International: Genetics. 53, 102493. doi:10.1016/j.fsigen.2021.102493

Wang, W.Y., Srivathsan, A., Foo, M., Yamane, S.K., Meier, R. 2018. Sorting specimen-rich invertebrate samples with cost-effective NGS barcodes: Validating a reverse workflow for specimen processing. Molecular Ecology Resources. 18, 490–501. doi:10.1111/1755-0998.12751

World Economic Forum. 2020. The Global Risks Report 2020. [https://www.weforum.org/reports/the-global-risks-report-2020].

Yeo, D., Srivathsan, A., Puniamoorthy, J., Maosheng, F., Grootaert, P., Chan, L., Guénard, B., Damken, C., Wahab, R.A., Yuchen, A., Meier, R. 2021. Mangroves are an overlooked hotspot of insect diversity despite low plant diversity. BMC Biology. 19. doi:10.1186/s12915-021-01088-z

Zhang, H., Bu, W. 2022. Exploring large-scale patterns of genetic variation in the COI Gene among Insecta: implications for DNA barcoding and threshold-based species delimitation studies. Insects. 13, 425. doi:10.3390/insects13050425

