## Supplementary Figures for "Rapid species discovery and identification with real-time barcoding facilitated by ONTbarcoder 2.0 and Oxford Nanopore R10.4"

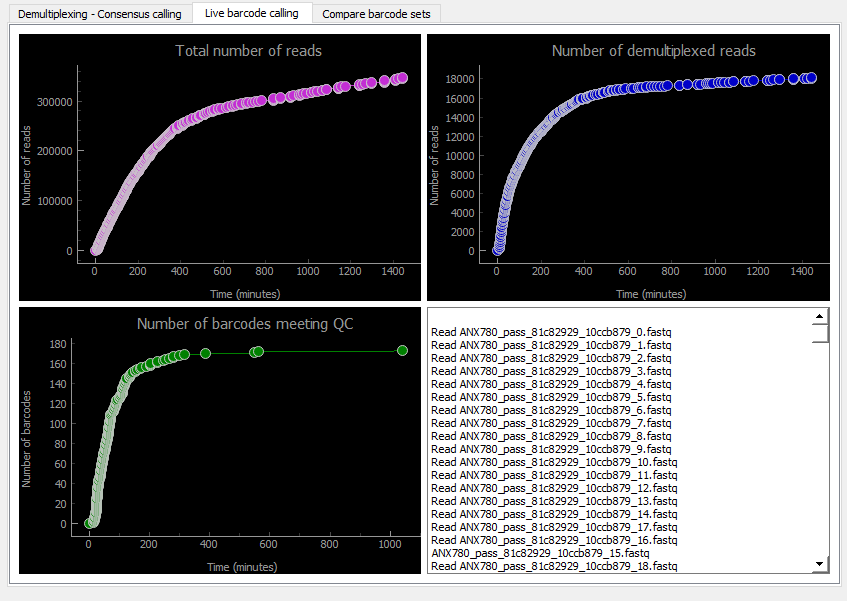

Supplementary Figure 1: The original real-time barcoding analysis conducted during the experiment involving Flongle R10.4 and the pool of 285 phorids: Top Left: Total number of reads base-called, Top Right: Number of reads demultiplexed used for barcoding calling, Bottom left: Number of barcodes called, and Bottom right: List of files processed.

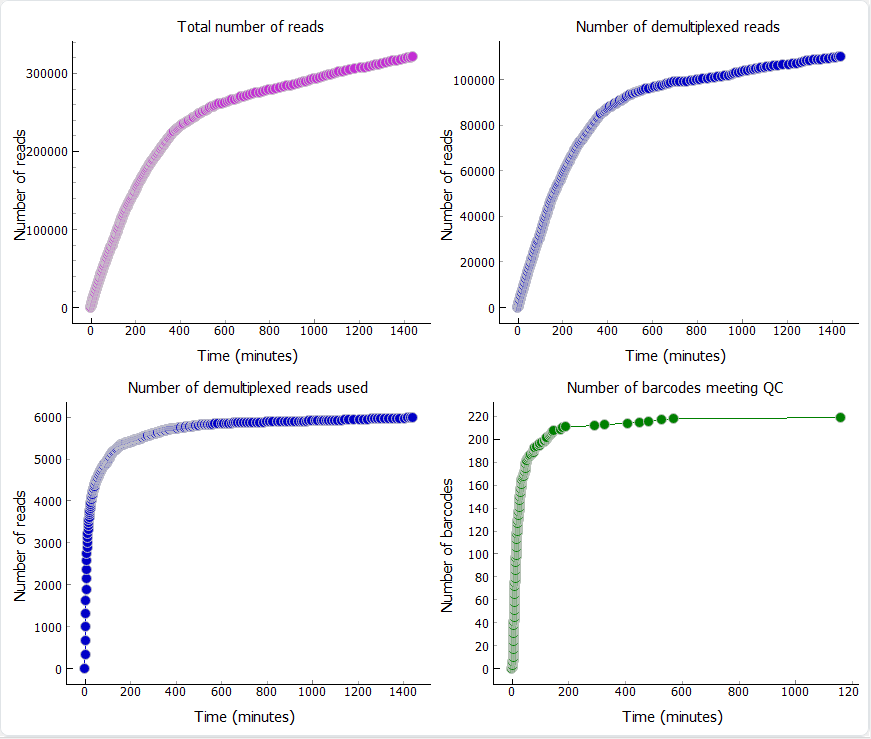

Supplementary Figure 2: Real-time barcoding based on reads base-called using high-accuracy model. The experiment involved Flongle R10.4 and a pool of 285 phorids.

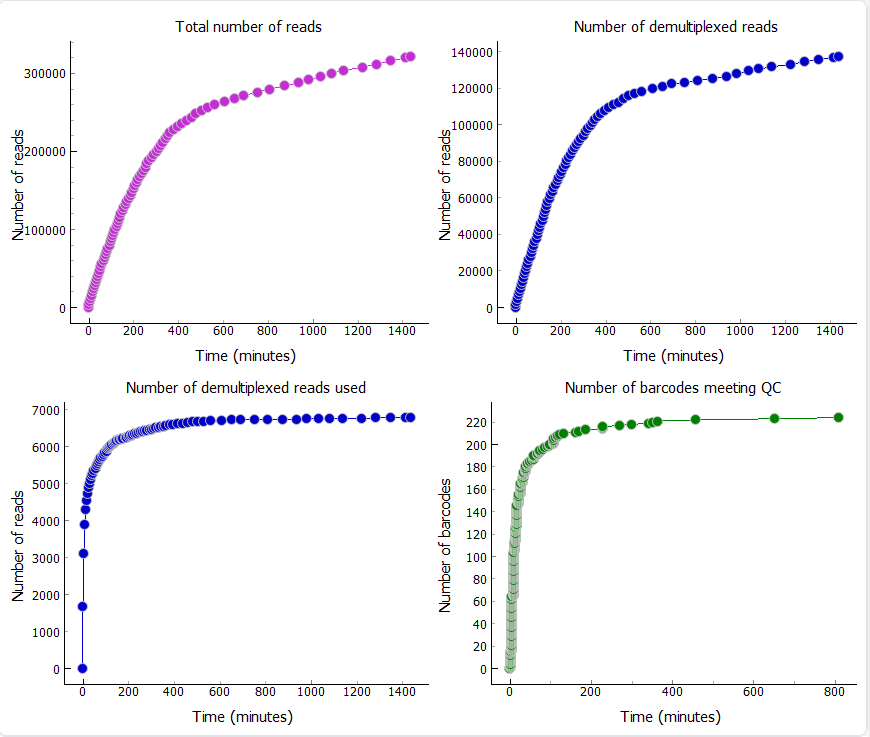

Supplementary Figure 3: Real-time barcoding based on reads base-called using super-accuracy model. The experiment involved Flongle R10.4 and a pool of 285 phorids.
